## Supplemental File for "Macrophage network dynamics depend on haptokinesis for optimal local surveillance"

### Supplemental Information

Supplemental Table

Legends to Supplemental Videos S1–S9

### Supplemental File 1

Supplemental Table for overview on all statistical tests performed.

| Figure | Statist. test | | Posthoc test | |
|---|---|---|---|---|
| 1D | **Kruskal-Wallis** | H=61.4, n(group)=3, P≤0.0001<br>N(Ctrl:CytoD)=25,25,<br>N(Ctrl:Y27)=25,25 | **Dunn's**<br><br>Control:CytoD<br>Control:Y27 | <br><br>*** (P≤0.001)<br>*** (P≤0.001) |
| 1E | **ANOVA** | F(2, 6)=172.0, P≤0.0001 | **Dunnett's**<br><br>Control:CytoD<br>Control:Y27 | <br><br>*** (P≤ 0.001)<br>*** (P≤0.001) |
| 1G | **Kruskal-Wallis** | H=70.62, n(group)=4, P≤0.0001<br>N(WT: $Tln1^{-/-}$)=25,25,<br>N(WT: $Itgb2^{-/-}$)=25,25,<br>N(WT: $Itgb1^{-/-}$)=25,25 | **Dunn's**<br><br>WT: $Tln1^{-/-}$<br>WT: $Itgb2^{-/-}$<br>WT: $Itgb1^{-/-}$ | <br><br>*** (P≤0.001)<br>ns (P>0.05)<br>*** (P≤0.001) |
| 1H | **ANOVA** | F(3,8)=58.43, P≤0.0001 | **Dunnett's**<br><br>WT: $Tln1^{-/-}$<br>WT: $Itgb2^{-/-}$<br>WT: $Itgb1^{-/-}$ | <br><br>*** (P≤0.001)<br>ns (P>0.05)<br>*** (P≤0.001) |
| 1 – fig suppl 1B | **unpaired two-tailed *t* test** | t=9.993, df=48, P≤ 0.0001 | - | *** (P≤0.001) |
| 1 – fig suppl 1E | **Mann Whitn. *U* test** | U=40, N(Ctrl)=25, N(CK-666)=55, P≤ 0.0001 | - | *** (P≤0.001) |
| 3D | **Kruskal-Wallis** | H=62.26, n(group)=3, P≤0.0001<br>N(Ctrl:CytoD)=25,25,<br>N(Ctrl:Y27)=25,25 | **Dunn's**<br><br>Control:CytoD<br>Control:Y27 | <br><br>*** (P≤0.001)<br>*** (P≤0.001) |
| 3E | **ANOVA** | F(2,6)=848.8, P≤0.0001 | **Dunnett's**<br><br>Control:CytoD<br>Control:Y27 | <br><br>*** (P≤0.001)<br>*** (P≤0.001) |
| 3F | **ANOVA** | F(3,96)=1.025, P=0.3851<br>N(WT: $Tln1^{-/-}$)=25,25,<br>N(WT: $Itgb2^{-/-}$)=25,25,<br>N(WT: $Itgb1^{-/-}$)=25,25 | **Dunnett's**<br><br>WT: $Tln1^{-/-}$<br>WT: $Itgb2^{-/-}$<br>WT: $Itgb1^{-/-}$ | <br><br>ns (P>0.05)<br>ns (P>0.05)<br>ns (P>0.05) |
| 3G | **ANOVA** | F(3,8)=0.5293, P=0.6746 | **Dunnett's**<br><br>WT: $Tln1^{-/-}$<br>WT: $Itgb2^{-/-}$<br>WT: $Itgb1^{-/-}$ | <br><br>ns (P>0.05)<br>ns (P>0.05)<br>ns (P>0.05) |
| 3 – fig suppl 1 | **unpaired two-tailed *t* test** | t=1.116, df=48, P=0.27 | - | ns (P>0.05) |
| 3 – fig suppl 2A | **ANOVA** | F(3,8)=2.926, P=0.0998 | **Dunnett's**<br><br>WT: $Tln1^{-/-}$<br>WT: $Itgb2^{-/-}$<br>WT: $Itgb1^{-/-}$ | <br><br>ns (P>0.05)<br>ns (P>0.05)<br>ns (P>0.05) |
| 3 – fig suppl 2C | **unpaired two-tailed *t* test** | t=6.643, df=48, P≤ 0.0001 | - | *** (P≤0.001) |
| 4D | **Mann Whitn. *U* test** | U=871, N(WT)=37, N($Itgb1^{-/-}$)=55, P=0.2455 | - | ns (P>0.05) |
| 5E | **ANOVA** | F(2,6)=418.8, P≤0.0001 | **Dunnett's** | |

| | | | | |
|---|---|---|---|---|
| | | | Control:CytoD<br>Control:Y27 | *** (P≤0.001)<br>ns (P>0.05) |
| 6C | **ANOVA** | F(3,8)=37.13, P≤0.0001 | **Dunnett's**<br><br>WT: Tln1$^{-/-}$<br>WT: Itgb2$^{-/-}$<br>WT: Itgb1$^{-/-}$ | <br><br>*** (P≤0.001)<br>ns (P>0.05)<br>*** (P≤0.001) |
| 6F | **ANOVA** | F(3,8)=33.30, P≤0.0001 | **Dunnett's**<br><br>WT: Itgb1$^{-/-}$ (2x) | <br><br>ns (P>0.05) |
| 6 – fig suppl 2 | **ANOVA** | F(3,8)=15.81, P≤0.001 | **Dunnett's**<br><br>WT: Tln1$^{-/-}$<br>WT: Itgb2$^{-/-}$<br>WT: Itgb1$^{-/-}$ | <br><br>ns (P>0.05)<br>** (P≤0.01)<br>ns (P>0.05) |
| 7D | **ANOVA** | F(3,8)=33.94, P≤0.0001 | **Dunnett's**<br><br>WT: Tln1$^{-/-}$<br>WT: Itgb2$^{-/-}$<br>WT: Itgb1$^{-/-}$ | <br><br>***(P≤0.001)<br>ns (P>0.05)<br>** (P≤0.01) |
| 7 – fig suppl 2 | **ANOVA** | F(3,8)=35.7, P≤0.0001 | **Dunnett's**<br><br>WT: Tln1$^{-/-}$<br>WT: Itgb2$^{-/-}$<br>WT: Itgb1$^{-/-}$ | <br><br>***(P≤0.001)<br>ns (P>0.05)<br>** (P≤0.01) |
| 8D | **unpaired two-tailed *t* test** | t=4.695, df=4, P=0.0093 | - | ** (P≤0.01) |

### Legends to Supplemental Videos S1 to S9

**Video S1: Random motility of macrophages in 3D matrices (related to Figure 1)**
Bone marrow derived macrophages (BMDMs) were cultivated from WT mice and embedded in matrigel. Live cell migration and shape changes were recorded over 20 h with phase-contrast microscopy using an Incucyte S3 instrument. This video relates to Figure 1B.

**Video S2: Talin-1 and β1 integrins control the mesenchymal shape and random motility of macrophages in matrigel (related to Figure 1)**
First part: Comparison of cell shape and migratory behavior of WT, *Tln1*$^{-/-}$, *Itgb2*$^{-/-}$ and *Itgb1*$^{-/-}$ BMDMs in matrigel. Live cell migration and shape changes were recorded over 20 h with phase-contrast microscopy using an Incucyte S3 instrument. Second part: Comparison of matrigel-embedded BMDMs from *Vav-iCre*$^{+/-}$ *Tln1*$^{fl/fl}$ *Lifeact-GFP*$^{+/-}$ and WT *Lifeact-GFP*$^{+/-}$ mice. Live cell shape changes and protrusion dynamics were recorded over 10 min with phase-contrast microscopy using confocal fluorescence microscopy. Lifeact-GFP signal is displayed as glow heatmap color. Third part: Comparison of cell shape and migratory behavior of WT, *Tln1*$^{-/-}$, *Itgb2*$^{-/-}$ and *Itgb1*$^{-/-}$ BMDMs in matrigel. Live cell migration and shape changes were recorded over 20 h with phase-contrast microscopy using an Incucyte S3 instrument. This video relates to Figures 1F–K and Figure 1 – figure supplement 1.

**Video S3: Integrin-independent chemotactic movement of macrophages in matrigel (related to Figure 3)**
First part: WT, *Tln1*$^{-/-}$, *Itgb2*$^{-/-}$ and *Itgb1*$^{-/-}$ BMDMs were embedded in matrigel and exposed to a chemotactic C5a gradient. Chemotactic migration along the gradient was recorded over 20 h with phase-contrast microscopy using an Incucyte S3 instrument. The lower row shows an animation of chemotactic cell tracks over the same time period. Second part: CK-666 treated and control BMDMs were embedded in matrigel and exposed to a chemotactic C5a gradient. Chemotactic migration along the gradient was recorded over 19 h with phase-contrast microscopy using an Incucyte S3 instrument. This video relates to Figures 3C, F and G and Figure 3 – figure supplement 1.

**Video S4: Amoeboid-like macrophages still perform chemotactic responses in mouse tissue (related to Figure 4)**
Two-photon intravital microscopy was performed on the ear skin of *Vav-iCre Itgb1*$^{fl/fl}$ *Lyz2*$^{GFP/+}$ and littermate control mice, which had received neutrophil-depleting Anti-Ly6G antibody to avoid the presence of neutrophils in imaging field of views. Chemotactic protrusion formation and cell body displacement toward a laser-induced tissue injury (orange area) of GFP-

expressing dermal macrophages in WT mice (first part) and conditional *Itgb1*-deficient mice (second part) were recorded over 90 min. GFP signal is displayed as glow heatmap color. This video relates to Figure 4.

**Video S5: Movement and protrusiveness as two sampling strategies for bead removal by macrophage networks (related to Figure 5)**

BMDMs were embedded in matrigel in the presence of fluorescent beads (red) coated with phosphatidylserine, an “eat-me” signal for macrophages. BMDMs were treated with vehicle or the ROCK inhibitor Y27632 and bead sampling activity recorded over 20 h with phase-contrast and fluorescence microscopy using an Incucyte S3 instrument. Control BMDMs are migratory and cover larger distance to sample beads (first part). Y27632-treated macrophages have reduced migration speed, but the increase in protrusiveness still allows efficient bead sampling (second part). This video relates to Figures 5C–H.

**Video S6: Haptokinesis is required for optimal bead removal by macrophage networks (related to Figure 6)**

β1 integrin-deficient BMDMs were embedded in matrigel in the presence of fluorescent beads (red) coated with phosphatidylserine, an “eat-me” signal for macrophages. Bead sampling activity was recorded over 19 h with phase-contrast and fluorescence microscopy using an Incucyte S3 instrument. Amoeboid-shaped *Itgb1*$^{-/-}$ BMDMs hardly move or form pronounced protrusions. Still, these macrophages are able to take up and ingest beads, but only beads in close vicinity to the cells. This video relates to Figures 6A–D.

**Video S7: Efficient efferocytosis of aged neutrophils by macrophage networks (related to Figure 7)**

WT BMDMs were embedded in matrigel in the presence of aged neutrophils that were labeled with pHrodo-Red (red). Fluorescent dead neutrophils were extracellular, before they were engulfed and removed by the macrophage network over time. Efficient efferocytosis in this assay relies on the interpretation of “find-me” and “eat-me” signals from dead neutrophils. The efferocytic activity of the macrophage network was recorded over 30 h with phase-contrast and fluorescence microscopy using an Incucyte S3 instrument. This video relates to Figures 7A and 7B.

**Video S8: Examples of efferocytic WT macrophages (related to Figure 7)**

WT BMDMs were embedded in matrigel in the presence of aged neutrophils that were labeled with pHrodo-Red (red). Three examples of macrophages that sequentially sample and ingest dead neutrophils are shown, revealing a phenotypic spectrum for efficient efferocytosis in WT cells. Cell 3 provides a good example of very directed protrusion formation toward dead neutrophil material. Efferocytic activity was recorded over 14, 28 and 18 h with phase-contrast and fluorescence microscopy using an Incucyte S3 instrument. This video relates to Figure 7E and Figure 7– figure supplement 2.

**Video S9: Haptokinesis is required for optimal efferocytosis by macrophage networks (related to Figure 7)**

WT and *Itgb1*$^{-/-}$ BMDMs were embedded in matrigel in the presence of aged neutrophils that were labeled with pHrodo-Red (red). WT BMDMs are migratory and cover larger distance to sample and engulf dead neutrophils. In contrast, *Itgb1*$^{-/-}$ BMDMs hardly move or form pronounced protrusions, allowing only uptake of dead neutrophils in close vicinity to them. Efferocytic activity was recorded over 23 h with phase-contrast and fluorescence microscopy using an Incucyte S3 instrument. This video relates to Figures 7E and 7F and Figure 7– figure supplement 3.
